# A RUNX2 GFP reporter is expressed prior to osteochondral differentiation and models Metaphyseal Dysplasia with Maxillary Hypoplasia and Brachydactyly (MDMHB)

**DOI:** 10.1101/2025.05.15.654100

**Authors:** Dimitrios V. Bikas, Sara Vardabasso, Gabrielle Quickstad, Karl B. Shpargel

## Abstract

SOX9 and RUNX2 are lineage defining transcription factors that drive differentiation of chondrocyte and osteoblast lineages respectively from osteochondral progenitors. In limb development, these progenitors are specified first by SOX9 expression required for mesenchymal stem cell (MSC) condensation prior to RUNX2 activation and osteochondral differentiation to chondrocyte and osteoblast lineages. Unlike limb development, the anterior craniofacial skeleton arises from cranial neural crest (cNCC) stem cells. To examine the temporal activation of SOX9 and RUNX2 within cNCCs, we utilized a combination of immunofluorescence to detect endogenous proteins and genetic reporters to label SOX9 and RUNX2 expressing cells. We find that RUNX2 is expressed broadly throughout cNCC stem cells of the first branchial arch that will give rise to developing mandibular tissue at a timepoint prior to osteochondral lineage determination. Substantial SOX9 expression is activated subsequently within differentiating chondrocytes. These findings were validated by fluorescent reporters inserted in the 3’ untranslated regions (3’UTRs) of *Sox9* and *Runx2*. Although the GFP based *Runx2* reporter did not delete any 3’UTR sequences, homozygous *Runx2^GFP/GFP^* pups develop postnatal deficiencies in intramembranous and endochondral ossification that correlate with enhanced expression of RUNX2 protein in osteoblasts and hypertrophic chondrocytes. *Runx2^GFP/GFP^*phenotypes model the human disorder, Metaphyseal Dysplasia with Maxillary Hypoplasia and Brachydactyly (MDMHB), resulting from RUNX2 enhanced activity due to intragenic duplications. Altogether, this reporter model provides a valuable tool for studying RUNX2 function in early cNCC-derived stem cell lineages and highlights the high sensitivity of ossification pathways to RUNX2 dosage.

**Summary:** We have developed a novel mouse model for a human disorder resulting from excessive RUNX2, a transcription factor required for bone formation. We find that RUNX2 turns on early within facial stem cells in a pattern unique from limb development. Excessive RUNX2 is particularly detrimental to bone growth in juvenile development after birth.

## Introduction

The mammalian skeleton arises from distinct lineages of progenitor cells whose specification is governed by master transcriptional regulators^1,2^. In vertebrates, ossification proceeds through either endochondral or intramembranous ossification, with the latter responsible for forming most craniofacial bones and clavicles. Intramembranous ossification advances through the direct differentiation of mesenchymal and neural crest progenitor cells into osteoblasts that remodel an extracellular matrix that becomes mineralized to form bone. In contrast, endochondral ossification is dependent on differentiation of progenitors to chondrocytes, a process which forms the majority of the skeletal system as well as the cranial base, providing structural support to influence facial morphology ^3–5^. Growth plate chondrocytes progress to a terminally differentiated hypertrophic state, secreting factors for extracellular matrix remodeling and mineralization, promoting vascular invasion, osteoblast recruitment, and bone formation. Both endochondral and intramembranous ossification require the RUNX2 transcription factor ^6,7^.

RUNX2 is a master transcription factor involved in osteoblast and terminal chondrocyte differentiation, processes that are essential for bone formation and growth ^6,8–10^. The RUNT domain of RUNX2 is the critical DNA-binding domain that regulates a wide range of bone-related genes. RUNX2 is considered a pioneer transcription factor, capable of accessing repressive chromatin to initiate accessibility and transcription ^2^. RUNX2 is required for commitment to osteoblast cell fate while Osterix (SP7) drives specification of osteoblast precursors ^6,7,11,12^. This process is tightly regulated by RUNX2 and Osterix ^13–15^. The observed presence of *Runx2* expression in Osterix knockout mice implicates RUNX2 as an upstream transcription factor in the regulation of osteoblast differentiation. No Osterix transcripts were detected in the skeletal elements of *Runx2* knockout mice, further validating Osterix as a downstream target of RUNX2 ^16^. Both *Runx2* and *Osterix* knockout mice are consequently devoid of osteoblasts and bone formation, emphasizing the necessity of both factors in skeletal development ^2,6,7,11,13,17^. RUNX2 deletion also leads to complete loss of chondrocyte hypertrophy critical for endochondral ossification ^7,15^. RUNX2 deletion specifically within chondrocyte lineages leads to reductions in proliferating chondrocytes with a loss of hypertrophic differentiation and extracellular matrix remodeling^15,15,18,19^. Although RUNX2 is required for subsequent hypertrophic differentiation, initial chondrocyte specification is dependent on the SOX9 transcription factor.

The appendicular skeleton and cranium originate largely from mesodermal mesenchymal stem cells (MSCs) and cranial neural crest stem cells (cNCCs) respectively ^20–24^. Limb endochondral ossification is dependent on MSC differentiation into chondrocytes and osteoblasts that form cartilage and bone. Reporter analyses has identified that both differentiated lineages develop from an osteochondral progenitor expressing the SOX9 transcription factor ^25^. The RUNX2 transcription factor is subsequently activated to drive osteoblast specification. SOX9 expression is maintained within chondrocyte lineages to repress RUNX2 and promote cartilage development ^26,27^. A *Sox9* knock-in mouse model with chondrocyte-specific *Sox9* overexpression resulted in a notable delay in the transition of proliferative to hypertrophic chondrocytes, accompanied by a similar stall in osteoblast differentiation ^28^. SOX9 is required within osteochondral progenitors as MSC specific mouse knockout fails to form both chondrocytes and osteoblasts leading to absence of limb cartilage and bone ^1^. In contrast, facial bones and osteoblast differentiation can develop normally with cNCC specific SOX9 mutation ^29^, highlighting distinct mechanisms regulating neural crest osteochondral development. RUNX2 is required within cNCCs to direct craniofacial intramembranous ossification, but the developmental timing of RUNX2 action is not well characterized ^14^.

Altered RUNX2 dosage is associated with human skeletal disorders ^6,9,30^. Haploinsufficiency of *RUNX2* is known to cause cleidocranial dysplasia (CCD), a rare autosomal-dominant skeletal disorder with high allelic heterogeneity and phenotypic variability, and is well-modeled in heterozygous loss of function mice ^6,30–32^. Clinical hallmarks of CCD include hypoplastic clavicles, cranial dysmorphism and dental anomalies. Metaphyseal dysplasia with maxillary hypoplasia and brachydactyly (MDMHB), an extremely rare skeletal disorder, results from heterozygous intragenic duplications of *RUNX2* concordant with a gain-of-function mechanism ^8,9,33–35^. The disorder is frequently characterized by widened clavicles, hypoplasia of cranial bones, short stature, and dystrophic teeth. Given its novelty and rare occurrence, MDMHB and its associated RUNX2 overexpression remain poorly modeled in current literature. RUNX2 has been overexpressed within osteoblast ^36,37^ or chondrocyte ^38^ lineages to disrupt endochondral ossification, however this method of experimentation results in excessive physiological levels of RUNX2 (greater than 8 fold enrichment compared to endogenous RUNX2) that prevent appropriate modeling of MDMHB.

We developed a GFP reporter to identify the timing of RUNX2 activation within cNCC lineages. While prior RUNX2 reporters using a distal promoter become activated during osteoblast maturation, we find that our reporter expressed from the 3′ untranslated region (3′UTR) of *Runx2* (*Runx2^GFP^*) becomes activated much earlier in cNCC lineages prior to osteochondral differentiation. These findings were validated by antibody detection of endogenous RUNX2 within broad regions of the first branchial arch that precede SOX9 upregulation in mechanisms distinct from limb MSCs. Although the reporter does not delete any 3’UTR sequences, homozygous *Runx2^GFP/GFP^* pups develop postnatal deficiencies in intramembranous and endochondral ossification that correlate with enhanced expression of RUNX2 protein. This reporter establishes an early cNCC model for RUNX2 function within stem cell lineages and validates that mechanisms regulating ossification are very sensitive to RUNX2 dosage.

## Materials and Methods

### Mice

The University of North Carolina Institutional Animal Care and Use Committee approved all animal research. The Runx2-Cre allele was generated using the distal promoter to drive CRE expression as described ^15^. *Sox9^GFP^* and *Rosa^Tomato^* reporter mice ^39,40^ were obtained from Jackson Labs (JAX stock # 030137 and #007909). *Runx2^GFP^* mice were developed by the University of North Carolina Animal Models Core. A cassette containing a trono spacer, internal ribosome entry site (IRES), Emerald GFP (EmGFP), and an FRT site were inserted immediately downstream of the Runx2 translation stop codon in exon 9 by CRISPR homology directed repair. The entire 3’ untranslated region (3’UTR) remained intact downstream of the cassette. The repair construct, enhanced specificity CAS9 protein, and gRNA (ATGGTTGACGAATTTCAATA) were microinjected into C57BL/6J embryos. Two independent founders were backcrossed to C57BL/6J to generate two independent lines of *Runx2^GFP^* that produce identical reporter properties and phenotypes. *Runx2^GFP^* lines were genotyped with long-range and vector backbone primers to ensure they have the correct insertion at the correct location.

### Embryonic Immunofluorescence

Tissue was fixed in 4% paraformaldehyde in 1X phosphate buffered saline (PBS) for 25 minutes (E11.5), 35 minutes (E12.5) or 40 minutes (E13.5) and processed for cryosectioning and immunofluorescence as described ^41,42^. RUNX2 (1:800, Cell Signaling 12556S or 1:150, Santa Cruz sc-390351), Osterix (1:150, Santa Cruz sc-393325), or Alexa-488 conjugated GFP antibody (BioLegend 338008) antibodies were incubated overnight at 4°C in blocking solution: 3%BSA, 10% goat serum, 0.1% Triton X-100, 1X PBS. Fluorescent conjugated secondary antibodies (Alexa-488 goat anti rat: Thermo-Fisher A-11006 or goat anti rabbit: A11008, Alexa-568 goat anti rabbit A-11036, or Alexa-647 conjugated goat ant mouse IgG1 A-21240) were incubated at 1:500 dilution for 1 hour at room temperature and mounted with ProLong Gold antifade (Thermo-Fisher P36930). Slides were imaged on a Leica fluorescent microscope or the Olympus VS200 slide scanner. Cellular fluorescence was calculated with Fiji/ImageJ2 using background subtracted integrated fluorescent density.

### Fluorescent reporter analyses and flow cytometry

Whole mount images of reporter fluorescence were captured with a Leica fluorescent stereomicroscope. The first branchial arch (BA) of E11.5 or the lower jaw of E13.5 embryos were dissected into PBS. E11.5 BA cells were dissociated as described ^43^ in cold protease solution (5 mg/ml of Bacillus Licheniformis protease: Sigma P5380, 5 mM CaCl2, and 125 U/ml DNase) at 4°C for a total 8 min, with trituration every 2 min using a P1000 pipet. Protease was inactivated with ice-cold RPMI containing 10% FBS and cells were passed through 40-micron strainer. E13.5 jaw tissue was dissociated as described ^44^. Briefly, jaw tissue was dissected, triturated with a P200 pipet, and was dissociated in HBSS containing 0.25% trypsin and 0.7mg/ml DNAseI for 10 minutes at 37°C with trituration after 5 minutes with a P1000 pipette. The trypsin was inactivated with 10% FBS and cells were passed through 40-micron strainer. Cells were subject to flow cytometry, flow sorting, or fixed with 2% PFA for 20 minutes at room temperature prior to nuclear staining as directed (BioLegend 424401) using RUNX2 (1:800, Cell Signaling 12556S) or Alexa-488 conjugated GFP antibody (BioLegend 338008). RNA from sorted cells was isolated (Trizol: Thermo-Fisher 15596026) and cDNA synthesized with ProtoScript II (NEB M0368L) prior to real time qPCR (Biorad 1725201) as described ^45^ for *Runx2*, *Sp7*, or *Gapdh*. Primers available upon request.

### Alizarin red and alcian blue staining

Samples were skinned and eviscerated. Samples were fixed in 95% EtOH for 2 days and stained in alcian blue (Amresco, 33864992) solution for 7-10 days. Alcian blue solution obtained by dissolving 0.15 g alcian blue in 800 mL 95% EtOH and 200 mL glacial acetic acid. Samples destained in 95% EtOH for 1 day, equilibrated with water (deionized) and transferred to 1% potassium hydroxide solution (KOH) for 1 day (P0 pups) or 3 days (P10 pups). Alizarin red S (Sigma, 130223) stock solution obtained by dissolving 0.25 g alizarin red in 1% KOH. Alizarin red S working solution obtained by diluting 250 μL alizarin red S stock solution in 100 mL KOH. Samples incubated in alizarin red S working solution for 1 day, rinsed with water (deionized) and transferred to glycerol for clearing. Images collected using Leica MZ FLIII Dissecting Scope with Nikon Digital Sight camera. Quantification of tibial, cranial base and mandible length conducted with QuPath *v0.4.3* software through *ROI Line: Annotations* tool.

### Postnatal fixation, decalcification, and paraffin embedding

Preparation process began with fixation in 4% paraformaldehyde/1x PBS (PFA, Electron Microscopy Sciences, 15710) for 2 days, decalcification in Immunocal (StatLab, 14141) for 3 days and dehydration in ethanol gradient at 30 minutes per stage (25%, 50%, 75%, 85%, 95%, 100%, 100%, 100% EtOH). Samples were passed through CitriSolv (3X, 30 minutes each, Fisher 22-143-975), 1:1 CitriSolv:paraffin solution for 60 minutes at 60°C and then incubated thrice in 100% paraffin for 30 minutes at 60°C. Samples changed to fresh paraffin and mounted in trays using Leica EG1160 Tissue Embedding Station.

### Antigen retrieval of paraffin sections and immunofluorescence

Paraffin embedding was performed as described Sections were mounted on slides, dried, and fixed in 2% PFA/1x PBS for 20 minutes and deparaffinized twice in CitriSolv for 5 minutes. Samples rehydrated in ethanol gradient at 2 minutes per stage (100%, 100%, 95%, 95%, 70%, 50%, 30% EtOH). Samples submerged twice in water (deionized) for 5 minutes and incubated overnight in diluted 1x ImmunoDNA (BioSB, BSB0020) solution at 55°C. Immunofluorescence was performed as described in Embryonic Immunofluorescence section.

### Hematoxylin and eosin staining

Hematoxylin (Harris Hematoxylin, Leica, NC0338284) filtered through cheesecloth before use. Eosin Y stock obtained by dissolving 2g Eosin Y (Fisher Scientific, 17372871) in 40 mL water, then adding 160 mL 95% EtOH. Eosin Y working solution obtained by diluting 100 mL Eosin Y stock in 300 mL 80% EtOH solution, then adding 2 mL glacial acetic acid. H&E staining was performed as described ^46^. Briefly, samples deparaffinized twice in CitriSolv for 5 minutes and rehydrated in ethanol gradient at 2 minutes per stage (100%, 100%, 95%, 95%, 70%, 50% EtOH). Samples submerged twice in 1x phosphate buffered solution (PBS) for 5 minutes. Samples stained in hematoxylin solution for 10 minutes, followed by 2 dips in water (deionized) then 2 dips in 2% glacial acetic acid solution. Samples dipped in 1.5% ammonium hydroxide 15 times followed by 10 dips in water and 1 minute stain in eosin Y working solution. Samples dehydrated in ethanol gradient at 2 minutes per stage (95%, 95%, 100%, 100% EtOH) and submerged twice in CitriSolv for 5 minutes. Samples mounted to glass coverslips with Citramount (Electron Microscopy Sciences, 1800501). Images collected using Leica DMi8 inverted microscope with Leica DFC9000 GT/GTC sCMOS camera. QuPath *v0.4.3* software used to measure growth plate thickness through *ROI Line: Annotations* tool and hypertrophic chondrocyte area through *ROI Polygon: Annotations* tool.

## Results

### RUNX2 is expressed early in cNCC stem cells prior to osteochondral differentiation

Previous studies on limb development have shown that mesenchymal stem cells (MSCs) exhibit high levels of SOX9 expression. Reporter analyses further revealed that these SOX9-expressing progenitors contribute to both osteoblast and chondrocyte lineages in the limb ^25^. RUNX2 becomes active within a defined domain of MSCs in the E10.5 limb bud, with its expression being spatially and quantitatively constrained by SOX9-mediated repression ^27^. These osteochondral progenitors, which co-express the antagonistic transcription factors SOX9 and RUNX2, possess the potential to differentiate into either osteoblasts (under RUNX2 influence) or chondrocytes (driven by SOX9: Fig. 1A).

**Figure 1:**
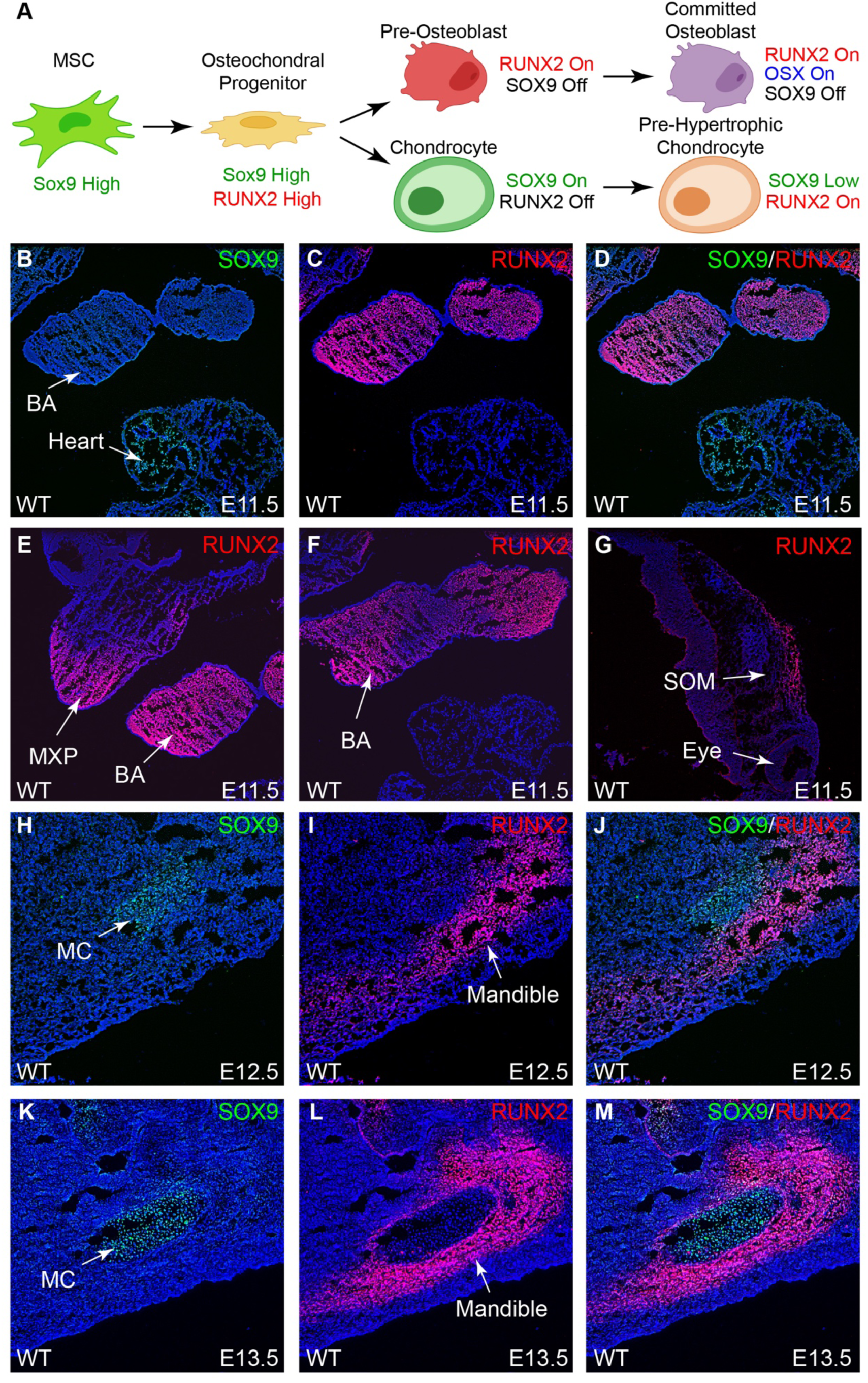
cNCC Runx2 is expressed prior to osteochondral differentiation. (A) Model for osteoblast and chondrocyte differentiation from MSC stem cells initially expressing SOX9 prior to RUNX2 induction in osteochondral differentiation. Model drawn with assistance of BioRender.com (B-E) IF for SOX9 (green) or RUNX2 (red) on coronal sections of E11.5 embryos overlayed with DAPI (blue). RUNX2 is expressed throughout the first branchial arch (BA) and proximal regions of the maxillary prominence (MXP) in the absence of SOX9 expression. (F) RUNX2 is expressed on the oral and proximal regions of more ventral BA sections. (G) RUNX2 is expressed in supraorbital mesenchyme (SOM) of the primordial frontal bone. (H-M) Mandibular SOX9 induction and osteochondral differentiation occurs across E12.5-E13.5 with SOX9 expression in chondrocytes of developing Meckel’s cartilage (MC) and RUNX2 expression in mandibular osteoblasts.

To investigate the temporal dynamics of SOX9 and RUNX2 expression in cranial neural crest cells (cNCCs), we performed immunofluorescence (IF) staining during early mandibular development in the first branchial arch (BA). RUNX2 expression was first detected in the BA mesenchyme at E11.5 (Fig. 1B-F), encompassing a broad population of cNCC-derived mesenchymal cells, despite only a subset ultimately committing to the osteoblast lineage. Interestingly, SOX9 was undetectable in the BA by IF at this stage, although robustly expressed in cardiac tissue, serving as a positive control (Fig. 1B). RUNX2 expression was highest in proximal and oral regions of BA sections (Fig. 1C), patterns that became accentuated in more dorsal BA domains (Fig. 1F). At this stage of development, RUNX2 was also active in maxillary prominences and supraorbital mesenchyme of the primordial frontal bone despite undetectable SOX9 in these regions (Fig. 1E and 1G). By E12.5 to E13.5, RUNX2 expression became more restricted to regions undergoing mandibular osteoblast differentiation, while SOX9 expression emerged in chondrocytes of Meckel’s cartilage (Fig 1H-M). These findings contrast with limb development, where SOX9 is predominantly expressed in osteochondral progenitors. In craniofacial development, RUNX2 is broadly expressed in cNCC-derived mesenchyme prior to overt osteochondral lineage segregation.

### SOX9 is expressed at reduced levels in BA cNCCs compared to limb MSCs

To enhance the sensitivity of detecting SOX9 expression in cranial neural crest cell (cNCC) lineages, we utilized a SOX9 fluorescent reporter line (*Sox9^GFP^*). This reporter was engineered to express EGFP via an internal ribosomal entry site (IRES) inserted into the 3′ untranslated region (3′UTR) of the *Sox9* gene (Fig. 2A), preserving endogenous gene function ^40^. Using this system, we readily observed EGFP fluorescence indicating SOX9 expression in the E11.5 limb bud. However, no detectable EGFP signal was observed in the facial region at this stage (Fig. 2B-C). To quantify SOX9 expression, we dissociated E11.5 limb and first branchial arch (BA) tissues and performed flow cytometry. This analysis revealed that a relatively small fraction (24%) of BA cells expressed the SOX9 reporter (Fig. 2F-G), compared to a significantly larger proportion (45%) in the limb bud, where reporter intensity was dramatically higher (Fig. 2J). By E13.5, EGFP fluorescence became evident in facial tissues (Fig. 2D-E), and flow cytometric analysis of the dissected lower jaw showed a marked increase in EGFP+ cells (58%) with heterogeneous SOX9 expression levels much more comparable to limb tissues (Fig. 2H-I, 2K).

**Figure 2:**
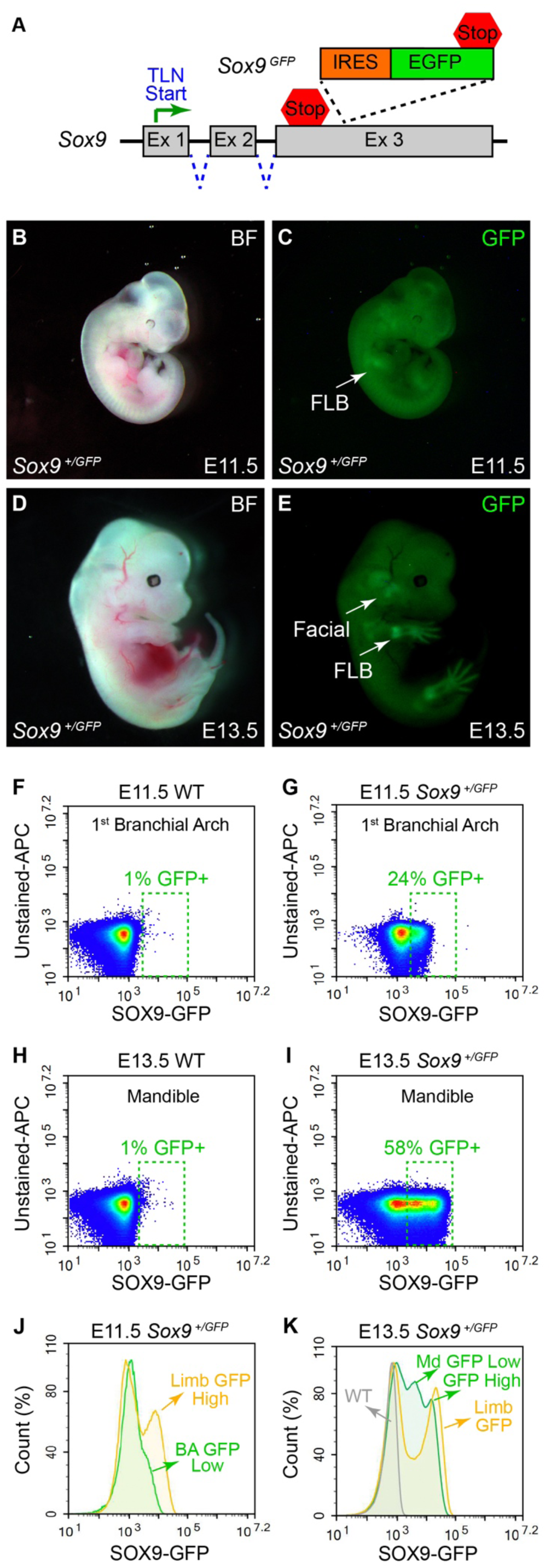
SOX9 is expressed at reduced levels in BA cNCCs compared to limb MSCs. (A) Model for *Sox9^GFP^* allele with insertion of an internal ribosome entry site (IRES) and enhanced green fluorescent protein (EGFP) reporter in the *Sox9* 3’ untranslated region (3’UTR). (B-C) Whole mount images of E11.5 *Sox9^+/GFP^* embryos detected GFP fluorescence within forelimb buds (FLB). (D-E) At E13.5, *Sox9^+/GFP^*embryonic fluorescence was detected throughout developing fore and hind limbs in addition to facial regions. (F-G) The E11.5 first BA of WT or *Sox9^+/GFP^* embryos was dissected and dissociated for flow cytometry. The X-axis of density plots illustrate average percentages of GFP+ cells from 3 biological replicates (boxed in green). (H-I) The E13.5 jaw was dissected for similar flow cytometry to parts F-G. *Sox9^+/GFP^* embryos display enhanced GFP+ percentages and levels of fluorescence compared to E11.5. (J-K) Histograms of flow cytometry in parts F-I with incorporation of limb cells for comparison. At E11.5, *Sox9^+/GFP^*levels are much higher in limb MSCs compared to BA cNCCs. At E13.5 as chondrocytes are specified, mandibular *Sox9^+/GFP^* levels experience heterogeneity with SOX9 levels more comparable to developing limbs.

### RUNX2 expression in cNCC stem cells is dynamic with subpopulation heterogeneity

To trace mandibular lineages originating from RUNX2-expressing cells in craniofacial tissues, we utilized a *Runx2-Cre* transgenic mouse line driven by the distal, bone-specific P1 promoter of *Runx2* (^15^, Fig. 3A). We crossed the *Runx2-Cre* transgenic line to *Rosa^Tomato^* mice, in which Cre-mediated recombination activates red fluorescent protein expression ^39^. Fluorescence analysis from E11.5 to E13.5 revealed no detectable *Runx2-Cre* activity at E11.5 (Fig. 3B-C); however, robust tomato fluorescence emerged at E12.5 and E13.5, first within the maxillary prominences (Fig. 3D-E) and later in mandibular and forelimb regions (Fig. 3F-G). Immunofluorescence at E13.5 confirmed Cre activity in only a subset of RUNX2-positive osteoblasts (Fig. 3N-P). Notably, tomato expression overlapped entirely with Osterix (Sp7) staining, indicating that the bone-specific *Runx2* promoter drives Cre expression exclusively in mature, committed osteoblasts (Fig. 3Q-R). Osteoblasts lacking Osterix expression (indicated by green arrows in Fig. 3N-R) did not exhibit reporter activity, further supporting the specificity of this Cre line for late-stage osteoblasts.

**Figure 3:**
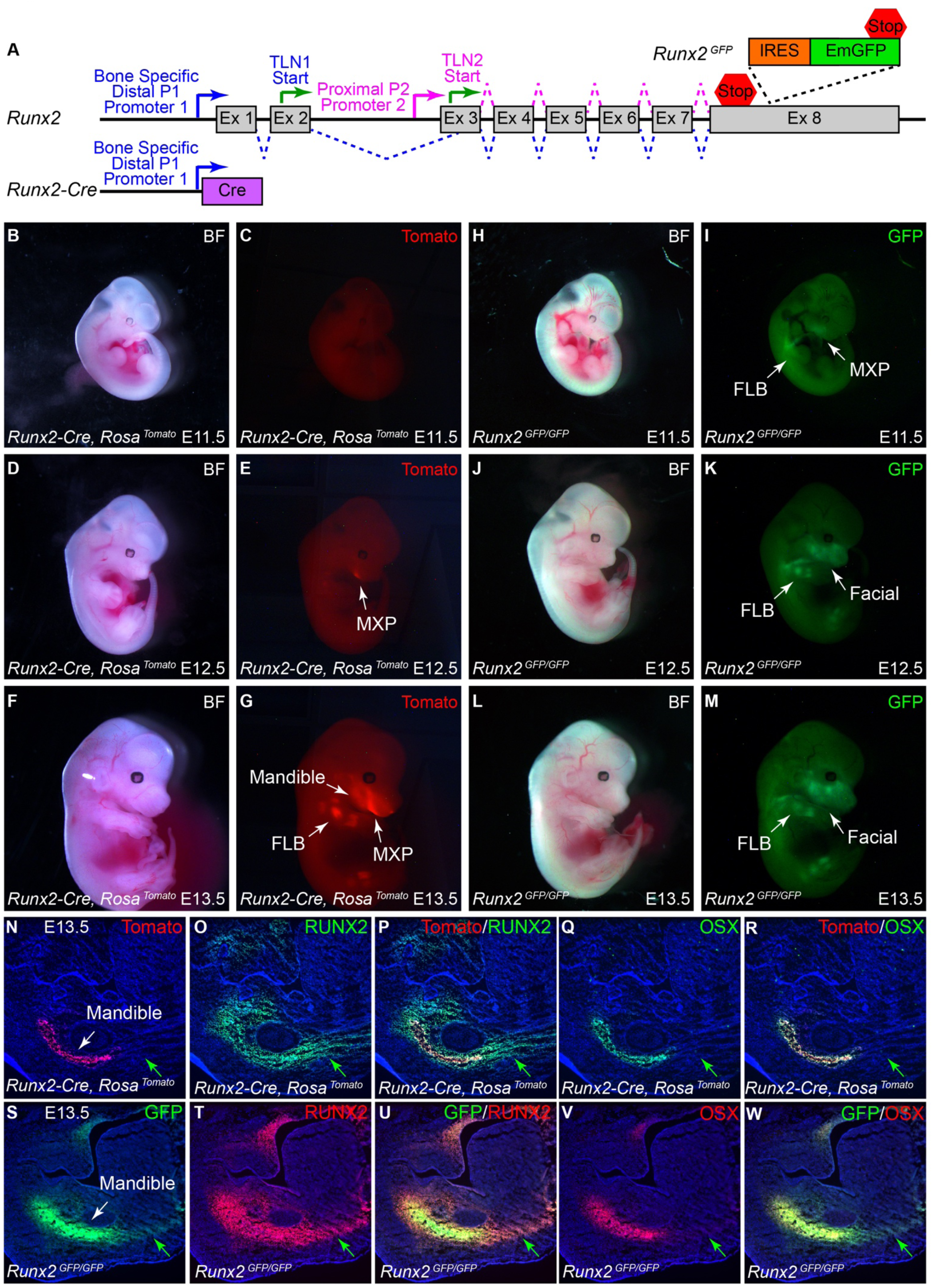
RUNX2 reporters label distinct populations of osteoblasts. (A) Schematic of both Runx2 reporters analyzed. The *Runx2-Cre* reporter places CRE expression under control of the bone specific distal promoter of *Runx2*. The *Runx2^GFP^* reporter inserts an internal ribosome entry site (IRES) and emerald GFP (EmGFP) reporter in the *Runx2* 3’ untranslated region (3’UTR). (B-G) A CRE activated *Rosa^Tomato^* allele was crossed with *Runx2-Cre* to label RUNX2+ cells by whole mount red fluorescence across E11.5-E13.5. *Runx2-Cre* induction of *Rosa^Tomato^*was absent at E11.5, initiated within maxillary prominences (MXP) at E12.5, and was more widely distributed in forelimbs (FLB), hindlimbs, maxillary, and mandibular regions at E13.5. (H-M) The *Runx2^GFP^* reporter exhibited green fluorescence at E11.5 in both forelimb bud and facial regions that increased and persisted across E12.5-E13.5. (N-R) E13.5 mandibular coronal cross sections of *Runx2-Cre* induced *Rosa^Tomato^* red fluorescence was overlaid with RUNX2 IF (green in O-P) or Osterix IF (OSX green in Q-R). *Runx2-Cre* induced *Rosa^Tomato^* fluorescence is specific for OSX+ mature osteoblasts but is not present in RUNX2+ OSX-pre-osteoblasts (green arrows). (S-W) E13.5 mandibular coronal sections of *Runx2^GFP^* reporter fluorescence (green) was overlaid with RUNX2 IF (red in T-U) or OSX IF (red in V-W). *Runx2^GFP^* reporter fluorescence is present in both pre-osteoblasts (green arrows) and mature OSX+ osteoblasts.

We hypothesized that early RUNX2 expression in E11.5 branchial arch (BA) cranial neural crest cells (cNCCs) may be driven by the proximal *Runx2* P2 promoter, which is active in a broader range of cell types, including non-skeletal lineages such as T cells. To more accurately reflect total *Runx2* expression, we generated a reporter line by inserting an IRES–emerald GFP (EmGFP) cassette into the 3′UTR of the endogenous *Runx2* gene (Fig. 3A). Given that our IF data indicate RUNX2 expression (Fig. 1) is initiated earlier than previously reported ^43,47^, this reporter provides a tool for validating those observations. In contrast to the *Runx2-Cre* system, which labels only committed osteoblasts, *Runx2^GFP/GFP^*embryos exhibited GFP fluorescence beginning at E11.5 in the maxillary and forelimb bud regions (Fig. 3H-I). This expression expanded to broader facial domains by E12.5–E13.5 (Fig. 3J-M). Because of low fluorescence sensitivity in *Runx2^GFP/GFP^* tissue sections, we used an anti-GFP antibody to more effectively detect labeled cells (Fig. 3S–W). This detection revealed complete overlap with RUNX2-expressing cells, including both pre-osteoblasts (indicated by green arrows in Fig. 3S–W, which lack Osterix expression) and mature, committed osteoblasts that co-express Osterix (Fig. 3V–W).

To validate reporter activity prior to osteochondral differentiation, we dissociated E11.5 BA tissue from *Runx2^GFP/GFP^* embryos and performed flow cytometry. Compared to wild-type (WT) controls, a substantial proportion of *Runx2^GFP/GFP^*BA cells (53%) were GFP-positive (Fig. 4A-B). Additionally, GFP-positive cells exhibited heterogeneous RUNX2 expression, with distinct populations expressing low or high RUNX2 levels (Fig. 4C). Co-staining with a RUNX2 antibody revealed that GFP expression from the reporter allele correlated proportionally with endogenous RUNX2 protein levels (Fig. 4D).

**Figure 4:**
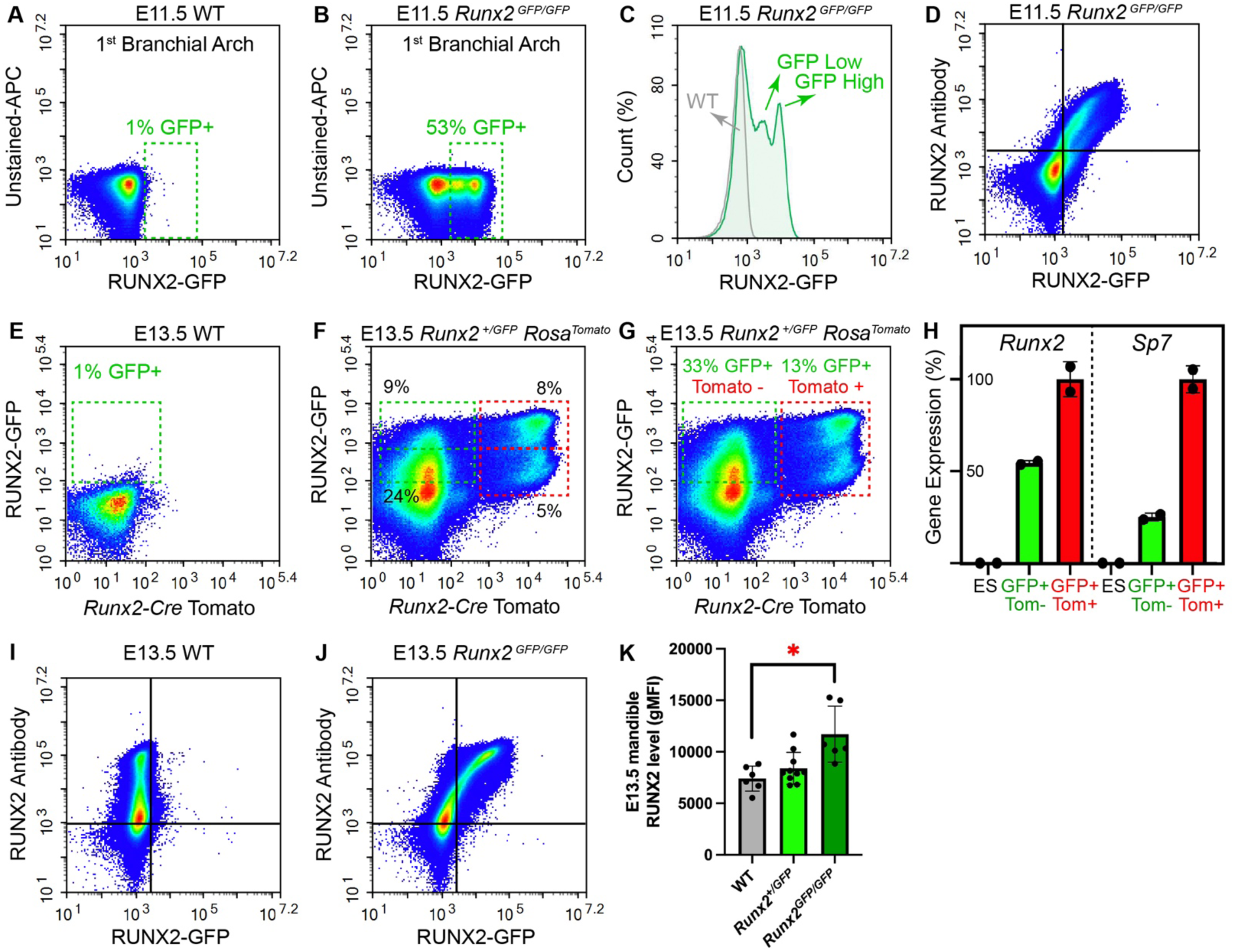
*Runx2^GFP^* fluorescence is present at E11.5 similar to IF results, demonstrates heterogeneity, and results in enhanced levels of endogenous RUNX2. (A-B) The E11.5 first BA of WT or *Runx2^GFP/GFP^* embryos was dissected and dissociated for flow cytometry. The X-axis of density plots illustrate average percentages of GFP+ cells from 3 biological replicates (boxed in green). (C) Histogram of flow cytometry in parts A-B identifies that RUNX2 is expressed at variable levels throughout BA cNCCs prior to osteochondral differentiation. (D) Nuclear staining with RUNX2 and GFP antibodies validated that GFP levels from the reporter correlate with endogenous RUNX2 protein. (E-F) The E13.5 jaws of WT or *Runx2^+/GFP^ Runx2-Cre Rosa^Tomato^* embryos were dissected for flow cytometry to contrast RUNX2 expression levels in pre-osteoblasts (Tomato-) or mature osteoblasts (Tomato+). *Runx2^+/GFP^* based fluorescence identified stratified low and high RUNX2 populations in both pre-osteoblasts (boxed green) and mature differentiated osteoblasts (boxed red). (G) Total populations of pre-osteoblasts and mature differentiated osteoblasts were sorted for RNA collection and cDNA synthesis. (H) qRT-PCR validated that *Runx2* and *Sp7* (Osterix) expression were enhanced in mature osteoblasts (red) compared to pre-osteoblasts (green). Mouse embryonic stem cell cDNA (ES) served as a negative control. Samples were normalized to *Gapdh* expression. (I-J) Nuclear staining with RUNX2 and GFP antibodies contrasts endogenous RUNX2 protein levels in WT or *Runx2^GFP/GFP^* dissociated mandibles. (K) Quantitation of geometric mean fluorescent intensity (gMFI) from WT, *Runx2^+/GFP^*, or *Runx2^GFP/GFP^* mandibles identified a significant increase in mean RUNX2 protein levels within homozygous *Runx2^GFP/GFP^* embryos.

To investigate RUNX2 protein dynamics during osteoblast differentiation, we combined the *Runx2-Cre Rosa^Tomato^* and *Runx2^GFP^* reporter alleles. Flow cytometry analysis of E13.5 *Runx2-Cre Rosa^+/Tomato^ Runx2^+/GFP^* embryos identified four distinct GFP-positive populations. Both tomato negative pre-osteoblasts and tomato positive mature committed osteoblasts displayed low and high levels of GFP expression (Fig. 4E-F), reflecting heterogeneity in RUNX2 levels. We isolated GFP+ tomato- (pre-osteoblasts) or GFP+ tomato+ (committed osteoblasts) populations by fluorescence-activated cell sorting (FACS, Fig. 4G), and qRT-PCR confirmed that the committed osteoblasts expressed higher levels of *Runx2* and *Sp7* (Fig. 4H). Overall, this dual-reporter strategy enables detailed dissection of RUNX2-regulated mechanisms underlying cNCC derived BA lineage specification and osteogenic differentiation during craniofacial development.

### *Runx2* 3’UTR insertion results in protein gain of function and disrupts postnatal bone growth

Co-staining with RUNX2 antibody identified that compared to WT, a greater proportion of E13.5 *Runx2^GFP/GFP^* mandibular osteoblasts express elevated levels of endogenous RUNX2 protein (Fig. 4I-K). This finding was unexpected, as the IRES–EmGFP insertion did not delete any native 3′ UTR sequences (Fig. 3A). We traced *Runx2^GFP/GFP^* development and pups were born (postnatal day 0 = P0) without any gross morphological abnormalities (Fig. 5A-C). Facial length was unaffected (Fig. 5D), however *Runx2^GFP/GFP^* pups exhibited a slight increase in facial angle (Fig. 5E) due to absence of curvature in the frontal-nasal junction (arrow in Fig. 5C). Additionally, bone and cartilage staining revealed no structural alterations to the mandible (Fig. 5F-G). As the mandible forms by intramembranous ossification, we also examined structure of the cranial base (Fig. 5H-I) and tibia (Fig. 5J-K) that form by endochondral ossification. All mandible, cranial base, and tibial bones analyzed demonstrated normal structure and measured dimensions in *Runx2^GFP/GFP^*pups at birth (Fig. 5L-O).

**Figure 5:**
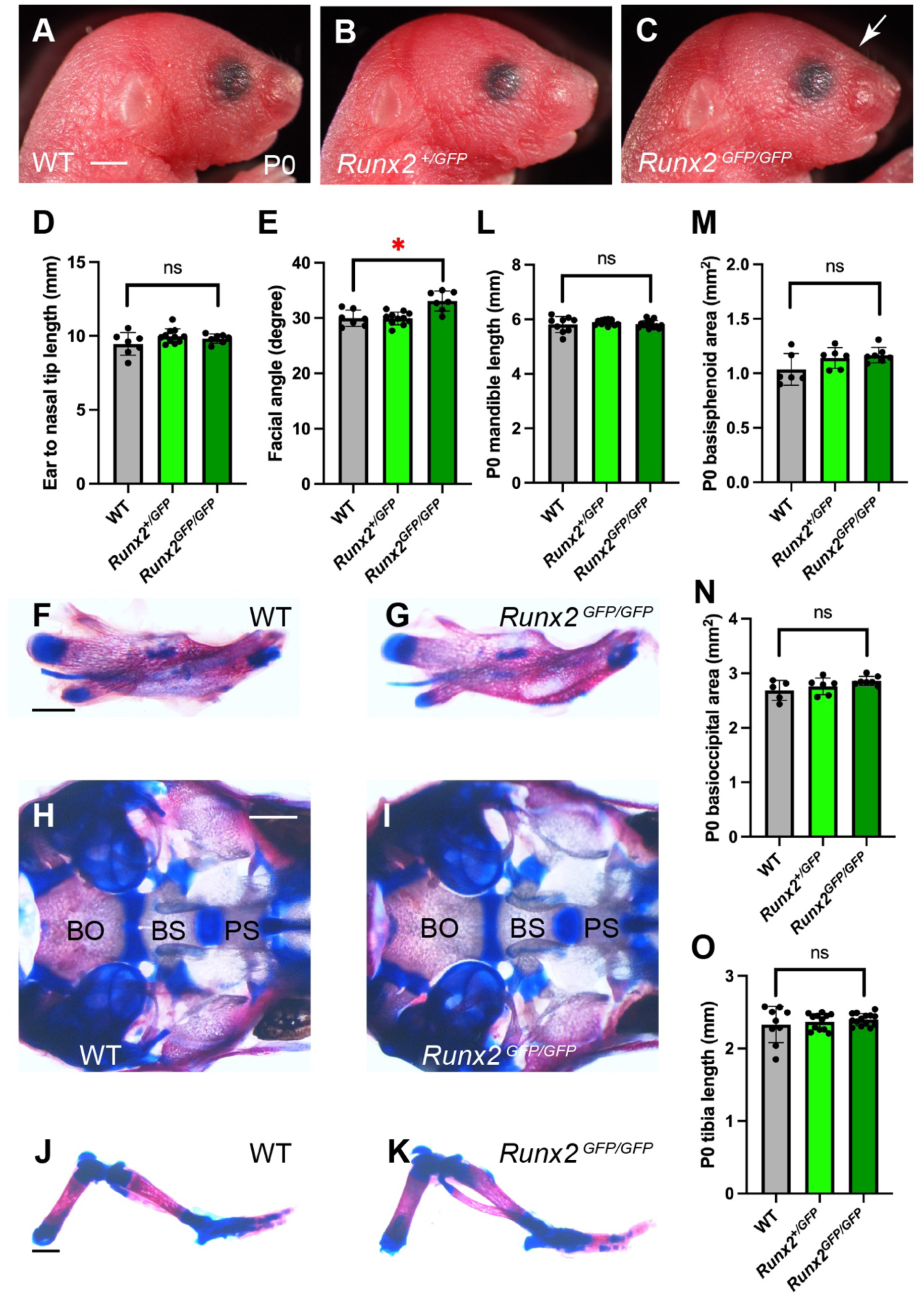
The *Runx2^GFP^* allele does not disrupt normal embryonic development. (A-C) Images of WT, *Runx2^+/GFP^*, or *Runx2^GFP/GFP^* pups at birth (postnatal day 0: P0). Although morphology similar *Runx2^GFP/GFP^* pups exhibited a slight absence of curvature in the frontal-nasal junction (white arrow). Scale bar = 2mm. (D-E) Quantitation of ear to nasal tip length revealed no abnormalities in P0 *Runx2^GFP/GFP^* pups despite a slight increase in facial angle due to forehead curvature. (F-G) Alizarin red and alcian blue skeletal stain of P0 mandible bone (red) and cartilage (blue) in WT or *Runx2^GFP/GFP^* pups. Scale bar = 1mm. (H-I) Ventral views of cranial base skeletal stain after mandible and hyoid removal. BO = basioccipital, BS = basisphenoid, and PS = presphenoid bones. Scale bar = 1mm. (J-K) Side views of skeletal stains of P0 hindlimbs. Scale bar = 1mm. (L-O) Quantitation of mandible length, basisphenoid area, basioccipital area, and tibia length revealed no abnormalities in the development of *Runx2^GFP/GFP^* bones at birth.

Although normal at birth, *Runx2^GFP/GFP^* pups were recovered in significantly reduced frequencies at weaning with fully penetrant malocclusion (Fig. 6A). Heterozygous *Runx2^+/GFP^*pups were observed at expected frequencies at weaning but did exhibit partial penetrance of malocclusion (Fig. 6A). *Runx2^GFP/GFP^* pups experienced growth deficiencies that manifested at P5 prior to malocclusion onset and persisted throughout juvenile development (Fig. 6B-F). *Runx2^+/GFP^* pups exhibited growth deficiency after P10 that tracked with the presence of malocclusion. *Runx2^GFP/GFP^*resulted in a smaller body and facial structures across P5-P6 (Fig. 6G-H) culminating in a dome shaped craniofacial structure at weaning (Fig. 6I). Alizarin red and alcian blue staining of P10 *Runx2^GFP/GFP^* bone and cartilage revealed shortened frontonasal structures (Fig. 6J-K, red arrow) and mandibles (Fig. 6L-M) indicative of disrupted RUNX2 function in osteoblast dependent intramembranous ossification.

**Figure 6:**
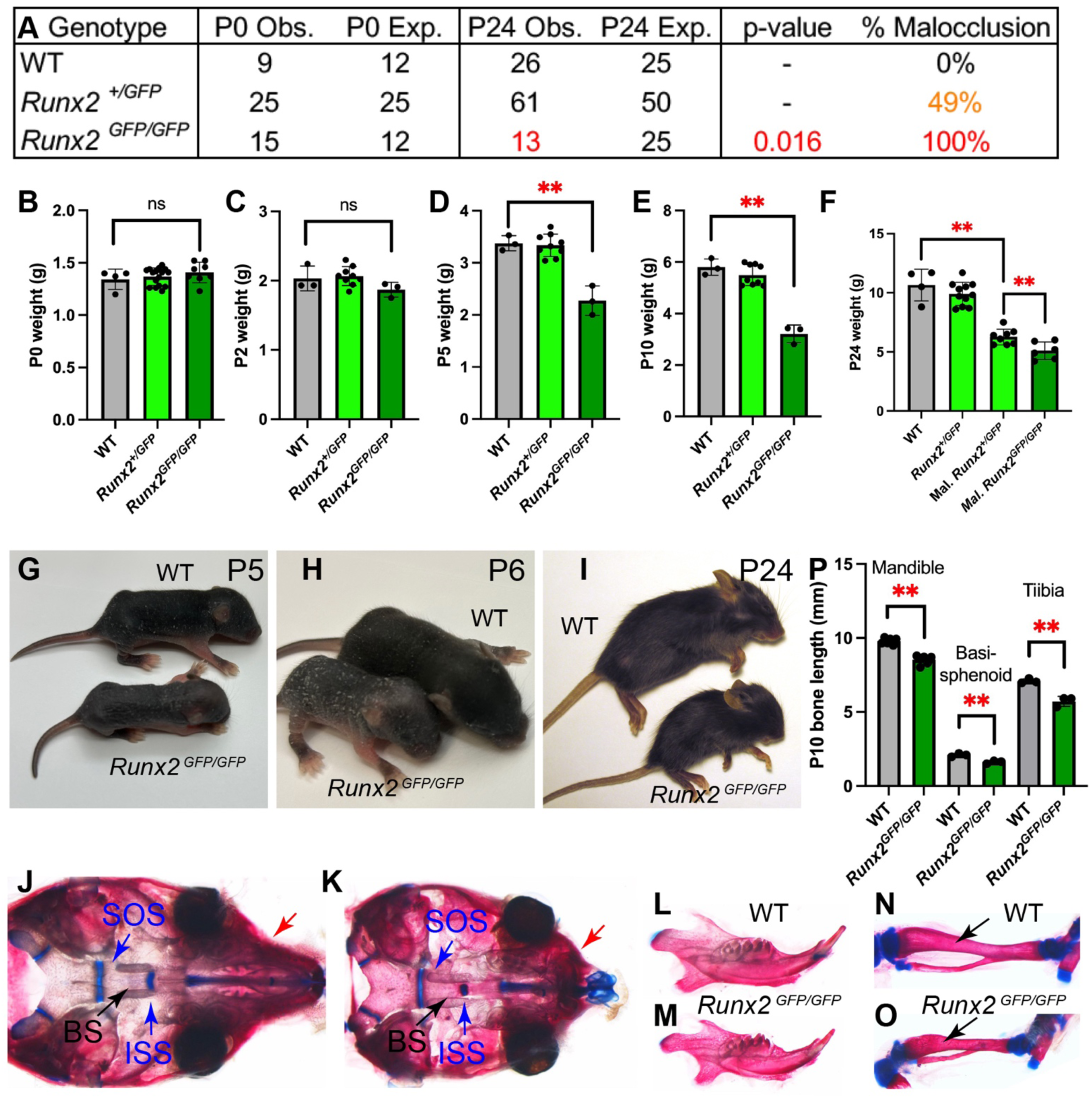
The *Runx2^GFP^* allele disrupts postnatal bone growth. (A) Observed frequencies of *Runx2* genotypes at birth (P0) and weaning (P24) and frequencies expected from heterozygous intercrosses. Red indicates observations underrepresented by chi-square p-value. Also illustrated us the malocclusion frequency. (B-F) Weights of *Runx2* genotypes between birth (P0) and weaning (P24). *Runx2^GFP/GFP^* pups experience growth deficiency starting at P5. *Runx2^+/GFP^* pups experience growth deficiency later after P10 due to malocclusion and difficulty feeding. (G-I) Images of *Runx2^GFP/GFP^* pups with comparison to WT at P5, P6, and P24. (J-O) P10 WT or *Runx2^GFP/GFP^* skulls and hindlimbs were stained for alizarin red and alcian blue to label bone and cartilage respectively. In J-K, the mandible was dissected, and images display the ventral view of the cranial base. BS = basisphenoid, ISS = intersphenoidal synchondrosis, SOS = spheno-occipital synchondrosis. *Runx2^GFP/GFP^* skeletal preparations displayed smaller frontal and nasal bone structures (red arrow) and partial closure of the ISS. *Runx2^GFP/GFP^* has smaller mandibles (L-M) and tibia (black arrows in N-O). (P) Quantitation of alizarin red stained lengths of indicated bones. *Runx2^GFP/GFP^* had significant reductions in mandible, basisphenoid, and tibia lengths. ** t-test p-value < 0.01.

Chondrocyte differentiation is essential for endochondral ossification, a process which forms the majority of the skeletal system as well as the cranial base, providing structural support to influence facial morphology ^3–5^. Growth plate chondrocytes activate RUNX2 to progress to a terminally differentiated hypertrophic state, secreting factors for extracellular matrix remodeling and mineralization, promoting vascular invasion, osteoblast recruitment, and bone formation. *Runx2^GFP/GFP^*pups also exhibited endochondral ossification deficiencies with shortened cranial base basisphenoid bones (Fig. 6J-K) and skeletal long bones (Fig. 6N-O) that develop from chondrocyte differentiation within growth plates. Quantitation of mandible, basisphenoid, and tibial lengths revealed moderate yet significant deficiencies in the growth of these bones (Fig. 6P). Notably, cranial base bones are developed by bidirectional growth from cartilage synchondroses. Basisphenoid formation is dependent on both spheno-occipital synchondrosis (SOS) and the intersphenoidal synchondrosis (ISS) cartilage. *Runx2^GFP/GFP^* skeletal preparations displayed a closing and loss of ISS cartilage (Fig. 6J-K). Therefore, enhanced RUNX2 expression disrupts normal intramembranous and endochondral ossification.

### RUNX2 upregulation impairs chondrocyte hypertrophic differentiation

We performed histology on P10 *Runx2^GFP/GFP^* pups to identify the cellular dysfunction leading to deficient postnatal endochondral ossification. Examination of tibial growth plates revealed that the columnar and hypertrophic zones of *Runx2^GFP/GFP^*chondrocytes had condensed (Fig. 7A-B) with a particularly significant reduction in hypertrophic thickness (Fig. 7I). Although *Runx2^GFP/GFP^* columnar chondrocytes were spaced regularly, these homozygotes experienced an increase in hypertrophic chondrocyte density (Fig. 7J), an indication that these cells may not be reaching a terminal differentiated state. *Runx2^GFP/GFP^*hypertrophic chondrocytes appeared smaller and more tightly packed (Fig. 7C-D), and calculation of cellular area identified that these cells were not expanding in size to a similar degree as WT chondrocytes (Fig. 7K). We examined cranial base synchondroses to identify if similar mechanisms were leading to basicranial bone growth deficiencies. At P10, the ISS located adjacent to basisphenoid and presphenoid bones was closing prematurely in the *Runx2^GFP/GFP^*basicranium (Fig. 7E-F). The *Runx2^GFP/GFP^* ISS bulged out of the cranial base with a unidirectional zone of hypertrophic chondrocytes that promoted fusion of the presphenoid and basisphenoid bones that normally does not occur in mice ^48^. The SOS that provides growth for basisphenoid and basioccipital bones was condensed in the *Runx2^GFP/GFP^*basicranium (Fig. 7G-H, 7I), similar to tibial growth plates. *Runx2^GFP/GFP^* SOS hypertrophic chondrocytes also demonstrated smaller cellular area indicative of incomplete terminal differentiation (Fig. 7K).

**Figure 7:**
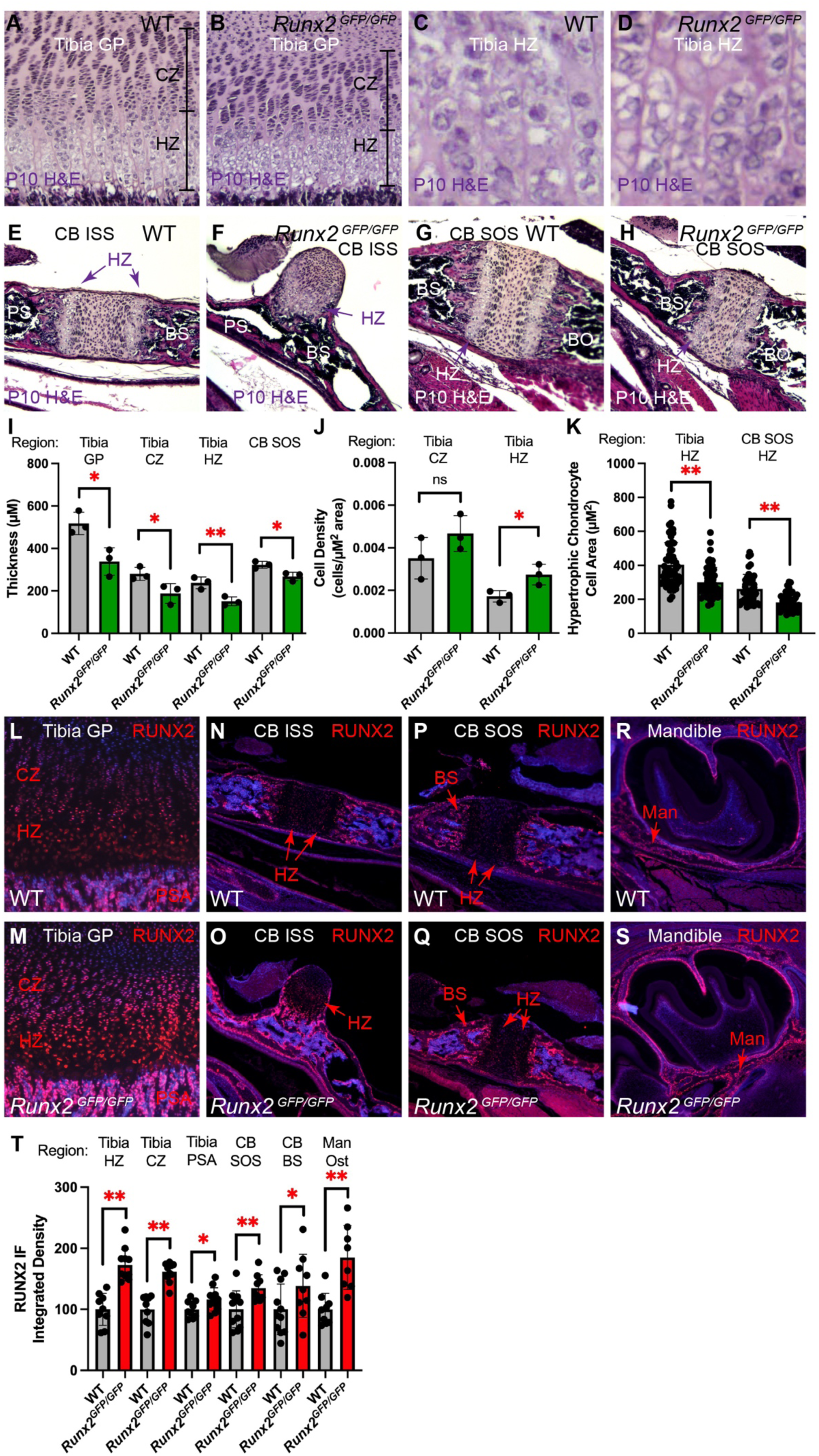
The *Runx2^GFP^* allele upregulates RUNX2 protein levels and alters chondrocyte differentiation at postnatal day 10. (A-B) H&E stained coronal P10 tibial sections illustrating proximal growth plate cartilage. Bracketed are chondrocyte columnar zones (CZ) and hypertrophic zones (HZ) based on cellular stacking and nuclear morphology. *Runx2^GFP/GFP^*tibias exhibit moderate reductions in columnar zone thickness and more dramatic reductions in hypertrophic zones. (C-D) Higher magnification view of tibial hypertrophic zone chondrocytes. *Runx2^GFP/GFP^* hypertrophic chondrocytes are smaller and more condensed. (E-H) H&E stained sagittal P10 midline cranial base sections of intersphenoidal synchondrosis (ISS in parts E-F) or spheno-occipital synchondrosis (SOS in parts G-H). Purple arrows denote hypertrophic zones of chondrocytes. PS = presphenoid bone, BS = basisphenoid bone, BO = basioccipital bone. *Runx2^GFP/GFP^* ISS is closing as PS and BS bones have fused and ISS has become a unidirectional adjacent bulge. *Runx2^GFP/GFP^* SOS is condensed with smaller thickness compared to WT. (I) Measurements of growth plate (GP), HZ, or CZ thickness in tibial sections or the cranial base (CB) SOS. Data points are individual pups with thickness calculated as an average from at least 3 measurements per growth plate across several sections. *Runx2^GFP/GFP^* experience condensed growth plates, synchondroses, and hypertrophic zones. (J) Calculation of cell density based on the number of columnar chondrocytes or hypertrophic chondrocytes per area of region analyzed. Data points are averages for individual pups with at 40 cells counted from several regions of interest per biological replicate. *Runx2^GFP/GFP^*hypertrophic chondrocytes have a higher cell density compared to WT. (K) Calculation of hypertrophic cell area. Data points are individual cell measurements from 3 biological replicates across several sections of tibial or SOS HZ. *Runx2^GFP/GFP^* hypertrophic chondrocytes have smaller area. (L-S) IF for RUNX2 (red) was overlayed with DAPI (blue) from P10 tibial growth plate, CB ISS or SOS, or mandibular bone (Man) sagittal cross sections of the molar region. RUNX2 protein in consistently upregulated in *Runx2^GFP/GFP^*chondrocytes (CZ and HZ) and osteoblasts in developing bone (PSA = primary spongiosa of the tibial, BS and Man). (T) Calculation of background subtracted RUNX2 IF fluorescent intensity (integrated density) based on several sections from 3 biological replicates. Values are normalized relative to WT (100%) for each region. RUNX2 protein is upregulated in all *Runx2^GFP/GFP^* chondrocytes and osteoblasts analyzed. Average increase across all cells and replicates is a 1.51-fold RUNX2 enhancement in *Runx2^GFP/GFP^*. * t-test p-value < 0.05 and ** t-test p-value < 0.01.

We performed IF for RUNX2 within various skeletal and craniofacial regions at P10 to identify if gain of function might be driving these cellular responses. RUNX2 protein was consistently elevated in tibial columnar chondrocytes, hypertrophic chondrocytes, and osteoblasts within the primary spongiosa (Fig. 7L-M). In cranial base synchondroses, RUNX2 was expressed in pre-hypertrophic chondrocytes prior to terminal differentiation (Fig. 7N-Q). RUNX2 was elevated in both *Runx2^GFP/GFP^*pre-hypertrophic chondrocytes and basisphenoid osteoblasts. Within the growing jaw, RUNX2 protein was increased in *Runx2^GFP/GFP^* mandibular osteoblasts (Fig. 7R-S). Quantitation of RUNX2 expression from all regions analyzed identified consistent RUNX2 enhancement within *Runx2^GFP/GFP^*chondrocytes and osteoblasts across multiple sections and biological replicates (Fig. 7T). The average enrichment across all sections analyzed was a 1.51-fold enhancement of RUNX2 protein within *Runx2^GFP/GFP^*chondrocytes and osteoblasts compared to WT samples. We conclude that appropriate RUNX2 dosage is critical for proper chondrocyte and osteoblast differentiation within the context of intramembranous and endochondral bone formation.

## Discussion

We now establish a mouse allele that upregulates RUNX2 protein as a model for the cellular and molecular consequences of RUNX2 gain of function in MDMHB. MDMHB results from intragenic duplication of RUNX2, within regions that minimally include exons 3-5 ^34^. This typically results in a RUNX2 protein with duplicated QA and Runt domains that mediate DNA binding and transcriptional activation. This modified protein encodes for approximately a 1.5-fold enhanced ability to transactivate RUNX2 target genes ^9^. We now demonstrate that mild upregulation (1.5-fold) of RUNX2 dosage in mice is capable of producing similar skeletal abnormalities including shortened skeletal long bones, facial and jaw hypoplasia, and dental anomalies. Although heterozygous *Runx2^+/GFP^*and homozygous *Runx2^GFP/GFP^* pups experience partially and fully penetrant malocclusion respectively, it is not clear if these phenotypes result from altered tooth development or inability to grind teeth due to altered jaw alignment from facial hypoplasia and abnormal cranial base structure.

RUNX2 exogenous overexpression was previously driven in chondrocyte lineages ^38^ resulting in deficiencies in bone growth. This system resulting from significant upregulation of RUNX2 (approximately 8-fold based on cell culture analyses) resulted in premature onset of chondrocyte hypertrophy in development. At perinatal timepoints, these embryos exhibited disorganized columnar and hypertrophic zones within growth plates at birth resulting in cellular apoptosis. The impact on perinatal chondrocyte differentiation was unclear as some hypertrophic markers appeared downregulated (*Col10a1*) while others were upregulated (*Mmp13*). We now establish a more moderate model of RUNX2 gain of function. *Runx2^GFP/GFP^* results in a 1.5-fold upregulation of RUNX2 protein, condenses columnar and hypertrophic zones, and restricts chondrocyte hypertrophy. The RUNX2 upregulation may accelerate transition from proliferative to hypertrophic chondrocyte states yet may prevent terminal differentiation. Other studies examining RUNX2 overexpression in osteoblast lineages ^36,37^ were also well beyond physiological levels (potentially 90-fold based on RNA levels ^37^). In these systems, bone development appeared unaffected ^37^, but transgenic mice experienced higher rates of bone resorption due to either enhancement of osteoclast activity ^37^ or impairment of osteoblast maturation ^36^. Our observation that early postnatal intramembranous ossification (frontal, nasal, mandible growth) is reduced in *Runx2^GFP/GFP^*pups indicates that mild RUNX2 upregulation from the endogenous locus is detrimental to osteoblast function. Future studies examining how enhanced levels of RUNX2 alters the transcription factor’s genomic distribution in osteoblasts and chondrocytes may shed more light on mechanism of altered dosage in skeletal disorder pathogenesis.

It is unclear why the 3’UTR insertion of IRES-EmGFP leads to enhancement of RUNX2 protein levels in *Runx2^GFP/GFP^* mice. The cassette is inserted immediately following the *Runx2* translation termination codon. The entire *Runx2* 3’UTR is left intact following the cassette. It is possible that a repressive regulatory site is present in the 3’UTR that becomes blocked by the insertion or expression of the of IRES-EmGFP cassette. *Runx2* expression is subject to 3’UTR repressive regulation by micro-RNAs ^49–53^. This insertion could also restrict other mechanisms repressing RUNX2 expression. *Runx2* is subject to m6A methylation by METTL3 within the 3’UTR that is read by YTHDF2 to promote mRNA degradation ^54^. There may be some other intrinsic positive regulatory information within the GFP cassette that promote more efficient transcription, RNA stability, RNA export, or translation. Additional experimentation examining specifically which regulatory mechanisms are supported by or lost due to 3’UTR insertion will clarify the responsible mechanisms and may lead to novel therapeutic strategies for CCD and MDMHB.

Through usage of the *Runx2-Cre* reporter, we find that the distal P1 bone specific promoter of *Runx2* is activated only in mature osteoblasts whereas other promoter regions must be driving *Runx2* expression in immature pre-osteoblasts and cNCCs within BAs and maxillary prominences. RNA *in situ* hybridization experiments have established that *Runx2-II* transcripts driven by the P1 bone specific promoter were present in more mature osteoblasts within developmental ossification centers but lacking in non-ossified mesenchyme and perichondrium ^55,56^. In contrast, *Runx2-I* transcripts expressed from the proximal P2 promoter were present throughout both regions. Loss of *Runx2-II* through deletion of the distal P1 promoter impaired endochondral ossification with more intact intramembranous ossification and experienced reduced expression of osteoblast maturation factors such as *Sp7* (Osterix) ^57^. Selective knockout of *Runx2-I* (stop codon after proximal P1 promoter) more severely disrupted intramembranous ossification compared to endochondral ossification with craniofacial development and cNCC originating tissue particularly affected ^58^.

We find that the *Runx2^GFP^* allele upregulates RUNX2 expression during both embryonic and postnatal development, yet *Runx2^GFP/GFP^* pups are both without obvious developmental abnormalities. It is not evident why postnatal development is particularly sensitive to excessive RUNX2 dosage. We find that RUNX2 is active early in embryonic development even prior to osteochondral differentiation in cNCCs of the first BA and maxillary prominences. These broad regions of expression may be essential for RUNX2 to regulate cNCC proliferation in a similar fashion to osteoblast lineages ^13^. In addition to osteoblasts, RUNX2 may play a role in establishment of alternative lineages derived from cNCCs including chondrocytes, fibroblasts, sensory ganglia, tendons, or surrounding connective tissue. At the stage of RUNX2 activation (E11.5), the first branchial arch begins to develop polarity with activation of proximal/distal and oral/aboral cell identities ^43,47^. RUNX2 expression appears particularly enriched within oral/proximal branchial arch locations and may play a more direct role in establishment of these lineages or driving regional specific proliferation impacting branchial arch morphogenesis. RUNX2 cNCCs specific knockout resulted in an absence of facial bone formation, but the early events in cNCC migration, proliferation, BA development, and establishment of cNCC lineages was not examined ^14^. Collectively, these data highlight several novel aspects of RUNX2 function for which future experimentation may provide additional information on pathogenesis and therapeutic targets for CCD and MDMHB.

## Funding

This research project was supported by NIH R01DE030530 awarded to K.B.S.

## Acknowledgements

Runx2-Cre mice were generously provided by Jan Tuckermann at the University of Ulm (Germany).

## Notes

### Competing Interest Statement

The authors have declared no competing interest.

